# Data-driven identification of major axes of functional variation in bacteria

**DOI:** 10.1101/2023.02.03.527046

**Authors:** Geneviève Lajoie, Steven W. Kembel

## Abstract

The discovery of major axes of correlated functional variation among species and habitats has revealed the fundamental trade-offs structuring both functional and taxonomic diversity in eukaryotes such as plants. Whether such functional axes exist in the bacterial realm and whether they could explain bacterial taxonomic turnover among ecosystems remains unknown. Here we use a data-driven approach to leverage global genomic and metagenomic datasets to reveal the existence of major axes of functional variation explaining both evolutionary differentiation within Bacteria and their ecological sorting across diverse habitats. We show that metagenomic variation among bacterial communities from various ecosystems is structured along a few axes of correlated functional pathways. Similar clusters of traits explained phylogenetic trait variation among >16,000 bacterial genomes, suggesting that functional turnover among bacterial communities from distinct habitats does not only result from the differential filtering of similar functions among communities, but also from phylogenetic correlations among these functions. Concordantly, functional pathways associated with trait clusters that were most important for defining functional turnover among bacterial communities were also those that had the highest phylogenetic signal in the bacterial genomic phylogeny. This study overall underlines the important role of evolutionary history in shaping contemporary distributions of bacteria across ecosystems.

**Originality-Significance Statement:** In this article, we use a trait screening approach based on genomic and metagenomic data to identify the key functional strategies of bacteria across ecosystems but also across the bacterial tree of life. This novel approach allows us to quantify the role of evolutionary processes in structuring microbial ecological differences among ecosystems. By reducing the high-dimensionality of trait variation observed among microorganisms around a small number of fundamental axes of trait covariation, we make a significant step towards generalization of the drivers of biological diversity in microbes but also across study systems. This research provide a major advance in our understanding of the origin and maintenance of bacterial biological diversity, expanding on related findings for plants and animals.

## Introduction

The identification of major axes of variation in life-history strategies among organisms and habitats has led to advances of both theory and practice in ecology and evolutionary biology (Grime 1977, Westoby et al. 1998, Lajoie and Kembel 2019). For example, in plants, large-scale screening of plant functional traits (Westoby 1998, Díaz et al. 2016) led to the discovery that a few major axes of covarying traits related to leaf economics, seed mass, and plant height explain most of the global variation in plant functional and life-history strategies. The contribution of this work has been twofold; first, it improved our understanding of the major environmental drivers and evolutionary origins of functional variation among organisms (Ackerly and Reich 1999, Wright et al. 2004). Second, the identification of a subset of measurable functional traits that could be used to characterize ecological strategies helped coordinate measurement efforts around a reduced set of interpretable functions, thus facilitating attempts at generalization among study systems worldwide (Díaz et al. 2016, Madani et al. 2018).

Recent conceptual and technological developments in the study of microbes have led to a proliferation of microbial trait measurements and screenings that have opened the door for extending this research agenda to the microbial realm. Still, functional categorization schemes proposed to date in microbial ecology have mostly consisted of direct translations of functional strategy schemes developed for macro-organisms. Classifications focusing on growth responses of organisms, such as the r-K selection spectrum or Grime’s CSR (competitive, stress-tolerant, ruderal) scheme (e.g. Fierer et al. 2007, Evans and Wallenstein 2014, Santillan et al. 2019), have drawn particular interest and been useful in improving our understanding of ecosystem functioning in certain habitats (Malik et al. 2020). The relevance of such schemes in explaining global patterns of microbial functional diversity however remains limited by their reliance on in-vitro phenotypic measurements, still unachievable for a vast diversity of microbial strains (Steen et al. 2019).

Metagenomic sequencing data represent a pragmatic alternative for the large-scale characterization of microbial functions. Functional predictions from metagenomic sequences have been useful for identifying sets of functional genes that best discriminate microbial samples across ecosystems (Dinsdale et al. 2008, Fierer et al. 2012, Ramírez-Flandes et al. 2019). For example, genes encoding oxidoreductases were shown to be the most useful of enzyme-encoding genes in separating metagenomic samples into biome groups (e.g. terrestrial vs. aquatic) (Ramírez-Flandes et al. 2019). By constraining their analyses along predetermined environmental axes, these studies allow the identification of functional genes that are distinctively enriched in each habitat in comparison to others. They however fall short in determining whether such ‘indicator genes’ take part in larger strategies that structure functional variation among microbial communities across diverse habitats – as observed for macro-organisms – or whether they tend to explain functional variation among habitat types idiosyncratically.

### Objectives and hypotheses

Here, we evaluate whether functional variation among bacterial metagenomes from different ecosystems is driven by major groups of correlated traits (i.e. functional strategies). We then compare these groups of traits to those explaining the most variation among a set of phylogenetically diverse bacterial genomes to test hypotheses about the ecological or evolutionary origin of these strategies. We predict that if the presence of correlated axes of trait variation across ecosystems is mostly explained by the filtering of groups of traits differentially selected along environmental gradients, then the groups of traits uncovered in genomes vs. metagenomes are unlikely to be the same between datasets (Appendix S1: Table S1). Reversely, if evolutionary selection or constraints on the variation of these groups of traits through time contribute to structuring functional variation among metagenomes (Futuyma 2010, Muir 2015), then the groups of traits should correspond among datasets. We lastly address the contribution of phylogenetic processes to correlated trait variation across metagenomes by testing if traits belonging to groups explaining the most variation among metagenomes also have stronger phylogenetic signal than other traits, and determine the phylogenetic depth at which they are most structured. Overall, this study contributes to a more general understanding of the role of functional strategies in driving bacterial distributions across ecosystems and of their evolutionary origins.

## Methods

### Metagenomic dataset collection and processing

We searched the IMG/M (Chen et al. 2019) and the MG-RAST (Keegan et al. 2016) databases for shotgun metagenomic sequence datasets from environmental and host-associated sources. We restricted our search to types of habitats for which we could obtain data from >1 study and for at least 5 different samples. To limit biases due to sequencing technology (Clooney et al. 2016), only datasets that had been sequenced with an Illumina system were included. To uniformize functional annotations among databases, we performed functional and taxonomic annotations of the IMG/M sequence datasets with the standard MG-RAST annotation pipeline (Keegan et al. 2016) using BLAST annotation identity criteria of a maximum e-value of 1e-10 and minimum percent identity of 30%, and a minimum sequence length of 125 nucleotides. To facilitate comparisons with the genomic dataset, metagenomic sequences were filtered to the bacterial domain only, which represented most annotated sequences (~94% of all taxonomic annotations). Past this filtering step, we only kept samples that had at least 100 000 sequences with both functional and taxonomic gene annotations, resulting in a dataset of 114 samples representing ten major bacterial habitats (Appendix S1: Table S2).

To control for differences in sequencing depth among samples, we randomly picked 100 000 sequences from each sample to generate functional and taxonomic community composition tables. Rarefying at that level was sufficient to capture the diversity of functional genes and taxonomic groups present in each sample (Appendix S2: Figure S1). To generate a functional composition table that had more interpretability and lower dimensionality, we summed counts of functional genes for each sample by functional pathway (i.e. Tier 3 KEGG category), the least aggregated of the three levels of the KEGG BRITE gene hierarchy (Kanehisa et al. 2014). Compositional data was transformed into relative abundance data for each sample. The resulting functional composition table spanned 245 functional pathways over 114 samples. A principal coordinates analysis of this dataset constrained by database of origin (i.e. IMG or MG-RAST) confirmed that database did not drive much variation among samples (1.9% of variation explained). The relative abundance of each taxon in each sample was similarly compiled at the phylum, class and order levels using the RefSeq taxonomic classification (Leary et al. 2016). The final taxonomic datasets spanned 28 phyla, 52 classes, 121 orders and 248 families of bacteria across all samples (Appendix S2: Figure S2).

### Genomic dataset collection and processing

The genomic dataset was retrieved from the AnnoTree server (accessed Dec. 2019) (Mendler et al. 2019). It consists of the functional annotations, taxonomic annotations and phylogenetic relationships of more than 27,000 fully sequenced bacterial genomes. We simplified the dataset by randomly retaining a single genome at each tip of the phylogeny, leading to a final dataset of 15,973 genomes (Appendix S1: Table S3). The taxonomic identity of all genomes retained spanned 111 phyla, 270 classes, 735 orders and 1659 families. The taxonomic composition of bacterial genomes in the genomic dataset encompassed the same major phyla recorded across metagenomes (Appendix S2: Figure S2). Counts of protein-coding genes for each genome were transformed into relative abundances of KEGG functional pathways as with the metagenomic dataset. We overall retrieved 312 pathways across genomes.

### Identification of main functional axes in the metagenomic dataset

We used a hierarchical clustering approach to identify groups of correlated functions that could constitute major ecological strategies across metagenomic samples. We first generated a principal coordinate analysis (PCoA) of the metagenomic functional dataset based on the Hellinger distance between samples using R package *ape* (Paradis and Schliep 2018). We then fitted the functional pathways onto the ordination to obtain vectors maximizing the correlation of these pathways with each ordination axis with R package *vegan* (Oksanen et al. 2013). To identify correlated axes of functional variation, we then calculated distances between functional pathways using the coordinates of these vectors across the first four PCoA axes (together explaining 74% of variance in the ordination). We used the resulting distance matrix to perform a hierarchical agglomerative clustering of functional pathways using Ward’s minimum variance method (D2) to minimize within-cluster variance. We identified clusters of correlated traits (i.e. trait clusters) by visual examination of the dendrogram based on their contribution to functional variation across the first 4 axes of the PCoA. The number of clusters in the dendrogram was confirmed based on a majority-ruling of 30 indices calculating the optimal number of clusters in a dendrogram using R package *NbClust* (Charrad et al. 2014). We then tested whether the number of pathways from each functional category differed among clusters when compared with their observed prevalence in the dataset using a chi-square test on counts of functional pathways by Tier 2 functional category in the KEGG classification framework, keeping only Tier 2 categories for which there were at least 10 observations across clusters.

### Drivers of functional variation among metagenomes

We evaluated if the richness of functional pathways within any sample varied among habitats using a type III analysis of variance controlling for unequal sample sizes among factors (Weisberg 2019). We compared habitat types in pairs using least square means comparisons using multivariate-*t* corrections with the R package *emmeans* (Lenth 2022). We then tested whether differences in functional richness among samples explained the functional distances among samples using a PERMANOVA. We also evaluated how much variation in the functional composition among samples could be explained by habitat type, taxonomic composition (class-level) and their interaction using a variation partitioning analysis. These analyses were performed using R package *vegan* (Oksanen et al. 2013).

### Phylogenetic structure of functional variation in the genomic dataset

We assessed the phylogenetic covariation of functional traits by performing a phylogenetic principal components analysis (pPCA) on the genomic functional dataset, using R package *adephylo* (Jombart and Dray 2010). This approach finds axes of functional variation that maximize the product of variance of the scores and their phylogenetic autocorrelation, revealing axes of correlated trait variation that are phylogenetically structured (Jombart et al. 2010). We calculated functional distances among pathways using the loadings obtained through this ordination over the first 4 axes of the pPCA, which encompassed ~65% of explained variance. We used these distances as described previously to perform a hierarchical clustering analysis of functional pathways using Ward’s minimum variance method.

We compared the structure of functional covariation in the metagenomic and genomic dataset with a Procrustes analysis performed on the cophenetic distance matrices describing each of the two hierarchical clusterings. The two dendrograms were pruned beforehand to include the same functional pathways. The significance of the Procrustes statistic, describing the similarity between the two datasets, was assessed by comparing the observed statistic to a distribution of 999 statistics generated through permutations of the original data (Oksanen et al. 2013). We generated a tanglegram of the two clusterings for visual comparison of their respective functional groupings. The trees were untangled prior to plotting using R package *dendextend* (Galili 2015).

### Phylogenetic signal and depth of ecologically important functions

We next evaluated whether pathways that drove functional differences among ecosystems tended to have high phylogenetic signal and to vary at more recent or ancient scales in the bacterial phylogeny. To test this, we calculated the phylogenetic signal of all functional pathways at increasing depths in the phylogeny using a phylogenetic correlogram. Confidence intervals were estimated through bootstrapping (n=100) of autocorrelation estimates after re-standardizations of phylogenetic weights (Keck et al. 2016). We then tested the effect of cluster affiliation, phylogenetic depth, and their interaction on the phylogenetic signal of traits using a Type III ANOVA for unbalanced designs. Differences among pairs of groups were tested at each depth using contrasts on estimated marginal means, as implemented in R package *emmeans* (Lenth 2022). We also tested whether the loadings of each pathway in the metagenomic PCoA was correlated with the strength of phylogenetic signal (absolute value of the signal) at each of 5 different depths using a linear model for each of the first four PCoA dimensions.

## Results

### Main strategies driving functional variation among metagenomes

Four major axes of functional variation explained nearly 75% of variance in the composition of functional pathways among metagenomic samples (Figure 1). These axes corresponded to functional transitions among major types of habitats, namely between animal guts, plants and soil, and aquatic habitats across the first 2 axes of variation (Figure 1ab). The third axis was characterized by functional variation between soil and aquatic sediment bacterial communities and saltwater ones, while the transition from (i) free-living and epiphytic habitats to (ii) endophytic ones characterized the fourth axis. Functional pathways could be divided into 9 main groups of correlated traits based on their contribution to functional variation across these axes (Figure 2a, Appendix S2: Figure S3, Appendix S1: Table S4). The proportions of functional categories to which the functional pathways belonged also differed significantly among groups (Tier-1 categories: *X*^2^(32, N=246) = 110.6, p<0.001; Tier-2 categories: *X*^2^(216, N=246) = 397.62, p<0.001) (Figure 2b).

**Figure 1.**
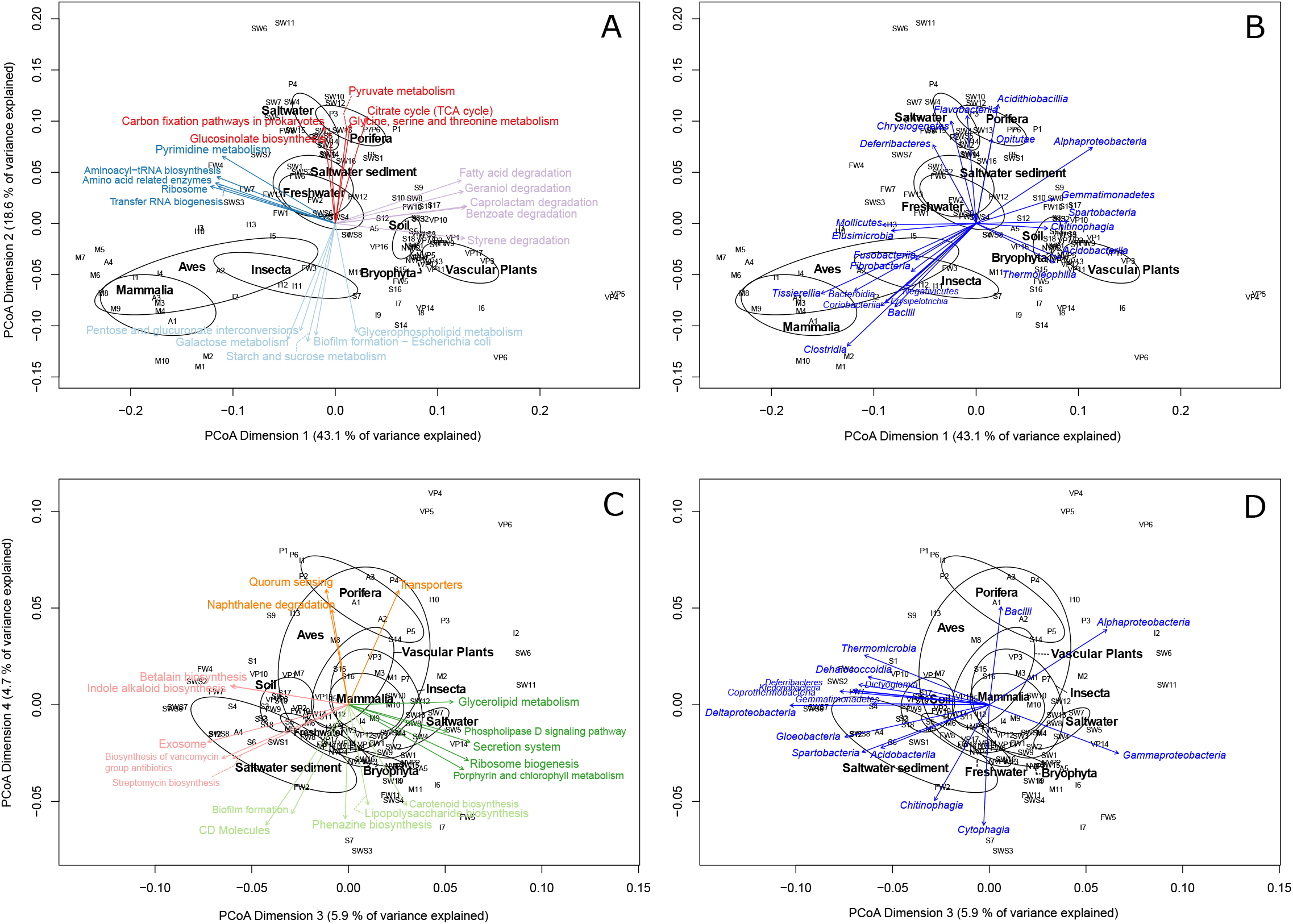
Principal coordinates analysis (PCoa) of bacterial functional pathways across habitats. Communities are represented by alphanumeric symbols where letters designate the habitat (A=Aves, B=Bryophyte, FW=Freshwater, I=Insecta, M=Mammalia, P=Porifera, S=Soil, SW=Saltwater, SWS=Saltwater sediment, VP=Vascular plants) and numbers designate the sample ID within that habitat (see Appendix S1: Table S2). Ellipses are drawn for each habitat and correspond to a 0.95 confidence interval from the group centroid. Functional pathways were fitted onto dimensions 1 and 2 (panel A), and dimensions 3 and 4 (panel C) of the ordination. Only the 20 functional pathways that were the most correlated with each set of dimensions are represented. They are colored by functional cluster (see Figure 2). Taxonomic classes composing these same communities were also fitted onto dimensions 1 and 2 (panel B) and dimensions 3 and 4 (panel D). All taxonomic classes whose relative abundances across samples were significantly correlated with each set of dimensions (α<0.001) are represented on the panels.

**Figure 2.**
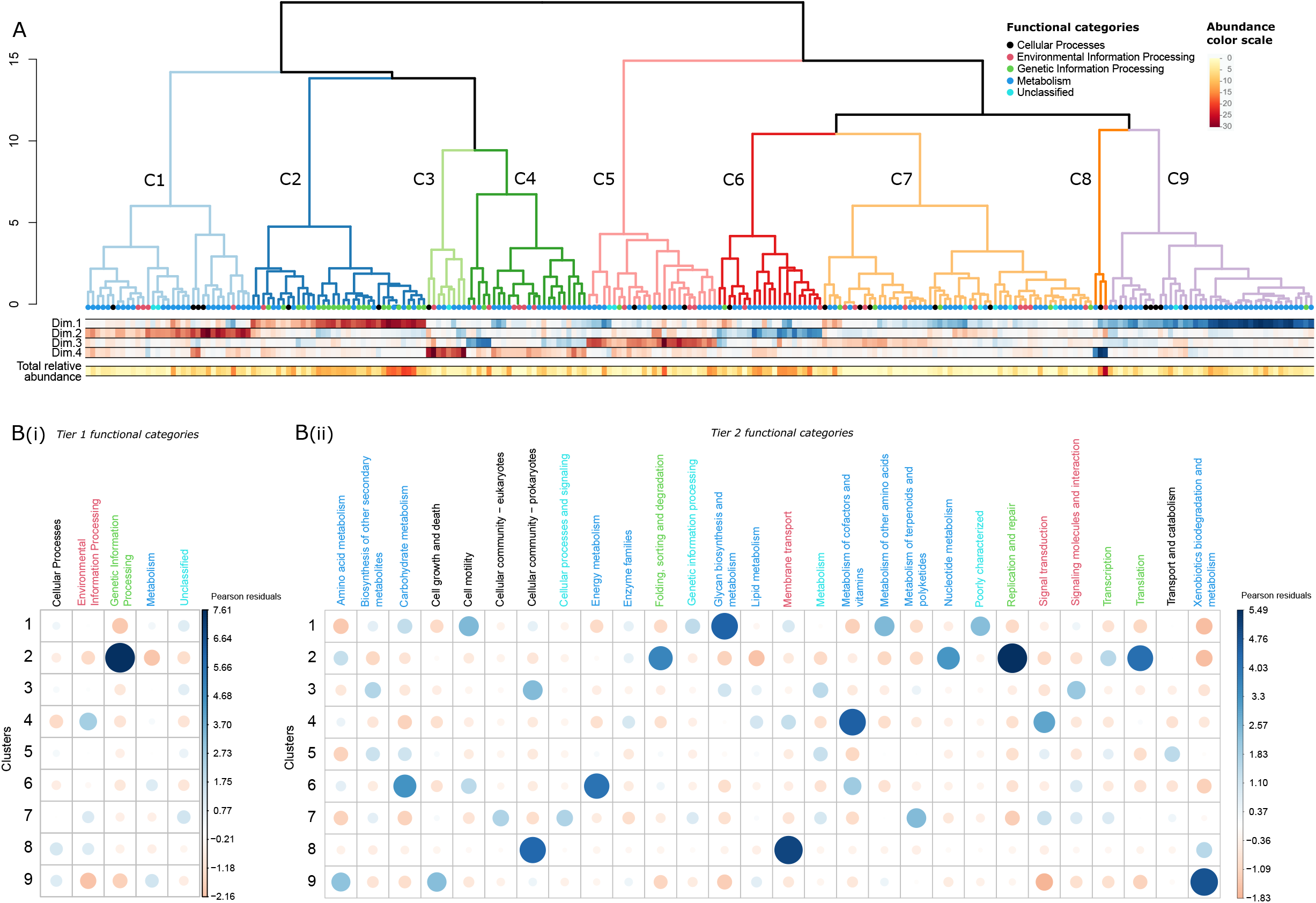
Hierarchical clustering of bacterial functional pathways across metagenomic samples (panel A) and contribution of functional pathways to the composition of each cluster, per Tier 1 and Tier 2 KEGG category (panel B). Panel A: Each dot at the end of branch represents a functional pathway, for which color codes for Tier 1 KEGG functional category (see legend). Each cluster (labelled C1 to C9) is color-coded to facilitate visual identification in other figures. The standardized explanatory power of each functional pathway across each of the first four dimensions of the metagenomic PCoA is indicated in a heatmap below the clustering. In the heatmap, blue is positively correlated with the dimension, and red is negatively correlated. The relative abundance of each pathway across all samples is also represented. Panel B: Dot size is proportional to the residuals of a chi-square test evaluating how different from normal is the distribution of functional pathways of a given functional category among clusters. The more functional pathways of a given category there is in a cluster compared to the expectation, the bigger and the bluer the dot (see legend). The less functional pathways of a given category there is in a cluster compared to the expectation, the bigger and the redder the dot. Color-coding of Tier-2 functional category follows that of panel A.

Habitat type explained an important portion of turnover in functional composition among samples (PERMANOVA: F(9,104) = 11.917, p=0.001, R^2^ = 51%). The functional composition of trait clusters can thus be described in relation to the habitats with which they are most associated. Namely, a cluster (C2) composed of traits pertaining to genetic information processing (i.e. replication and repair; translation; folding sorting and degradation) and nucleotide metabolism explained the functional differentiation of animal gut samples as compared with soil and plants samples (Axis 1). A cluster (C9) characterized by traits related to xenobiotics biodegradation and metabolism, amino acid metabolism as well as cell growth and death explained the variation in the opposite direction. Along the second axis, functional pathways (C6) participating in carbohydrate metabolism, energy metabolism (i.e. photosynthesis) and the metabolism of cofactors and vitamins were together enriched in samples from aquatic communities, while pathways (C1) participating in glycan biosynthesis and metabolism, metabolism of other amino acids and cell motility were enriched in terrestrial ones (both free-living and host-associated).

A cluster of traits (C4) involved predominantly in the metabolism of cofactors and vitamins and in environmental information processing (i.e. signal transduction) explained variation along the third axis and were enriched in the saltwater communities as opposed to the soil one. No clear predominance of any functional group emerges for the cluster (C5) correlated with the opposite side of this axis. Compared to other clusters, those (C3, C8) driving variation along the 4^th^ axis both contained pathways involved in prokaryote community sensing and cell-cell interactions. In contrast with cluster C8 associated with endophytic Porifera and plant microbiomes, the cluster (C3) correlated with non-endophytic communities was also enriched in functions related to the biosynthesis of other secondary metabolites.

While functional differences among samples could be linked to the turnover in the identity of pathways that characterize these communities, differences in the richness of functional pathways among samples explained 14% of variation in functional distances among samples in a constrained ordination. Indeed, the functional diversity of individual samples varied among bacterial habitats, as revealed for both richness (Type-III ANOVA: F(9,104) = 7.855, p<0.001) and Shannon diversity (Type-III ANOVA: F(9,104) = 7.695, p<0.001). Bacterial communities of animal guts (e.g. mammal, bird and insect guts) were generally the least functionally diverse while communities associated with plants (e.g. vascular plants and bryophytes) and soil were the most functionally diverse (Appendix S2: Figure S4). Controlling for the effect of richness on the PCoA however did not affect to a large extent the configuration of samples in the ordination (m12 squared = 0.5071, corr. = 0.7020, sig. = 0.001, # permutations = 999). Most variation in the functional composition of samples among habitats could be explained by turnover in the taxonomic composition of these communities (Figure 1bd, Appendix S2: Figure S5).

### Phylogenetic structure of correlated trait variation

The first four axes of the phylogenetic PCA explained 65% of phylogenetic variation in the genomic functional dataset. Clusters of functional pathways that were important in explaining functional variation among ecosystems were also important in explaining phylogenetic variation in the functional composition of genomes (m12 squared = 0.3618, corr. = 0.7989, sig. = 0.001, # permutations = 999) (Figure 3). Across all traits, the intensity of phylogenetic signal varied significantly as a function of depth, with higher signal at lower depths and lowest signal at intermediate depths. While the signal was positive at lowest depths, it became negative about mid-way towards the root of the tree. The strength of phylogenetic signal also varied significantly among metagenomic functional clusters and in interaction with phylogenetic depth (Figure 4, Appendix S1: Table S5). Clusters 2 and 9 exhibited more phylogenetic signal than other clusters at low phylogenetic depths, while clusters 1,5 and 7 exhibited the least. The latter clusters were also the least phylogenetically structured at deeper depths, while cluster 6 and cluster 2 were the most structured. These differences can be linked with variation in the mean phylogenetic depths of traits belonging to different Tier 1 category, with genetic information processing pathways (enriched in Cluster 2) showing the most signal across phylogenetic depths (Appendix S2: Figure S6). Functional pathways that contributed most to variation among metagenomic samples along the first PCoA axis (characterized by a trade-off between clusters 2 and 9) also consistently displayed the strongest signal in the genomic phylogeny (Appendix S2: Figure S7, Appendix S1: Table S6). This relationship was observed across most depths but was stronger at low phylogenetic depths. It was weaker or null across the other PCoA dimensions.

**Figure 3.**
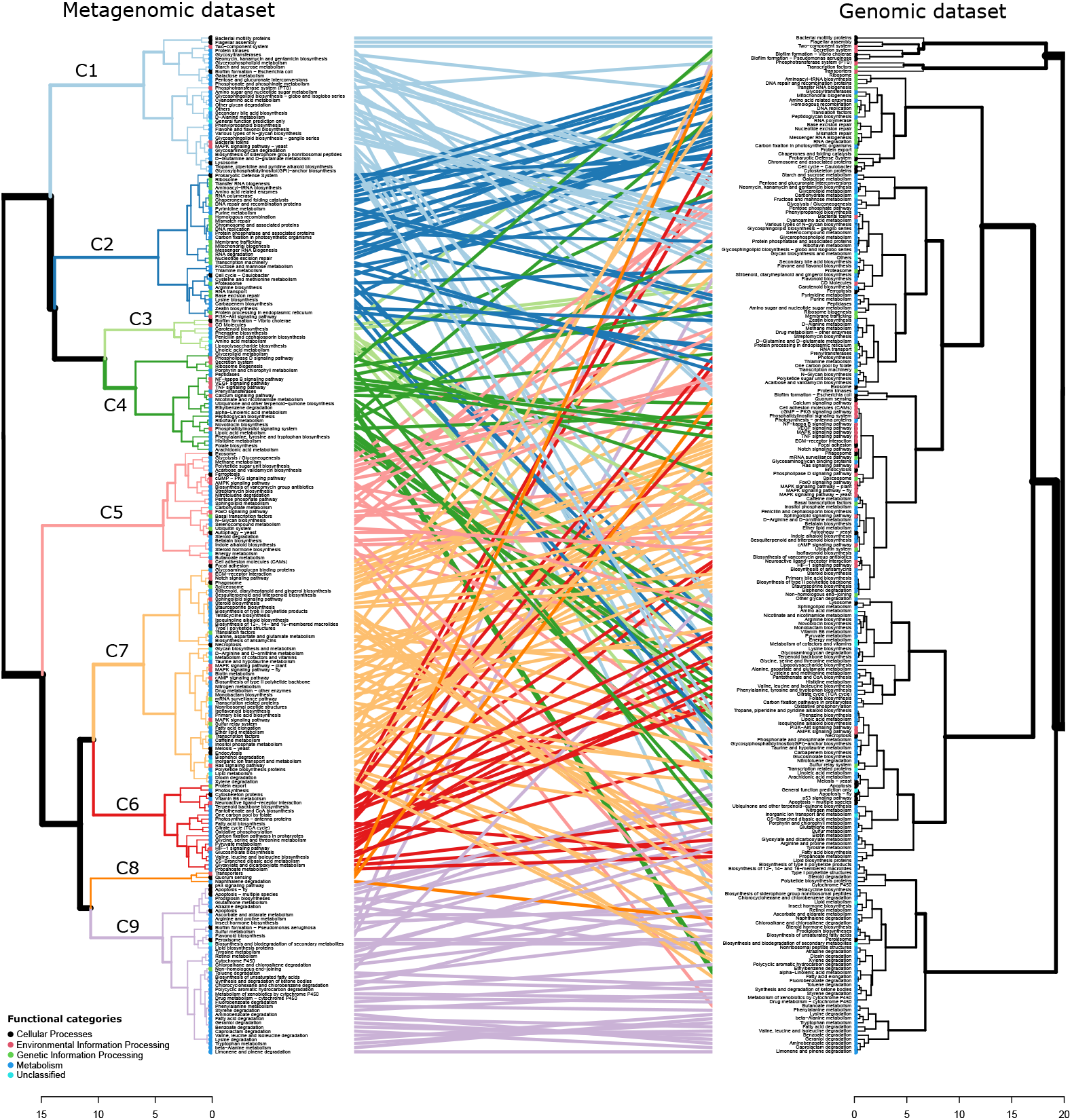
Tanglegram comparing functional clusters calculated from correlations between bacterial functional pathways across (left) metagenomic samples from different habitats and across (right) bacterial genomes. Hierarchical clustering of the genomic data is phylogenetically constrained (see Methods). Dots at the end of each branch are color-coded by the Tier-1 KEGG functional category to which the functional pathway belongs. Lines connect the same functional pathways among datasets and are color-coded by metagenomic cluster.

**Figure 4.**
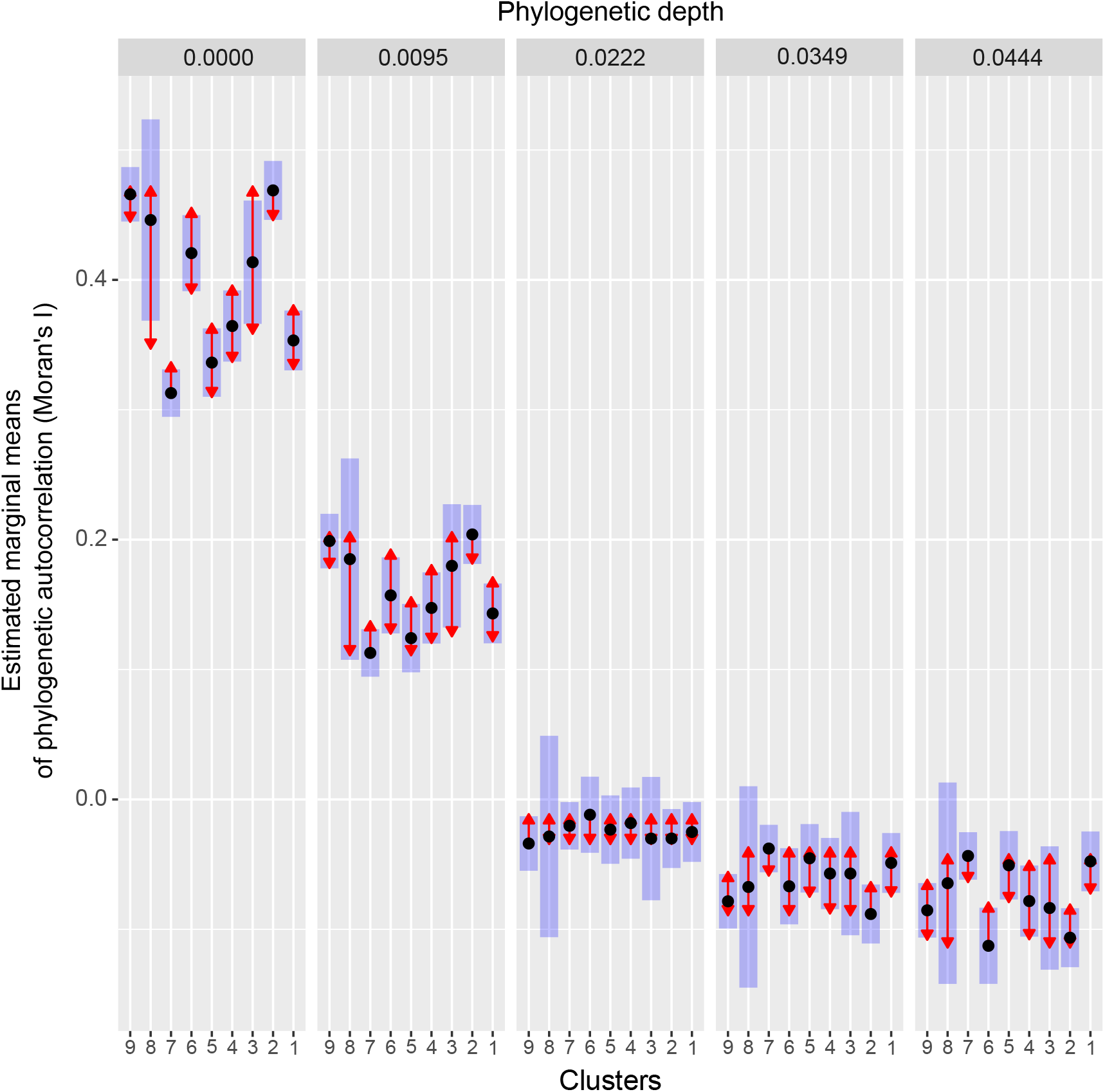
Phylogenetic signal (Moran’s *I*) of functional pathways among clusters and across phylogenetic depths. Moran’s *I* was calculated per pathway using phylogenetic correlograms across the bacterial genomic phylogeny, from the tip (depth = 0) towards the root of the tree (depth = 0.0444). Black dots represent the value of estimated marginal means of Moran’s I across the functional pathways of each cluster at each depth. The blue bars are confidence intervals. The absence of overlap among red arrows of two clusters at a given depth indicates that their mean phylogenetic signals are significantly different. These pairwise comparisons were adjusted for multiple testing using the multivariate *t* distribution method (Genz et al. 2021).

## Discussion

In this study, we show that functional variation among bacterial communities sampled across diverse habitats is structured by clusters of correlated functions defining contrasting life-history strategies among major bacterial habitat types. The structure of these clusters derived from metagenomic annotation data was similar to that obtained from the clustering of functional pathways among bacterial clades using a genomic dataset. This concordance suggests that functional turnover among bacterial communities from distinct habitats does not only result from the differential filtering of similar functions among these communities, but also from phylogenetic correlations among these functions. Functional pathways associated with trait clusters that were most important for defining functional turnover among bacterial communities were also those that were the most phylogenetically structured, underlining the important role of evolutionary history in shaping contemporary distributions of bacteria across ecosystems.

### Ecological significance of functional clusters characterizing contrasting habitat types

A few main clusters of correlated functional pathways explained functional variation among bacterial communities in our metagenomic datasets. The prevalence of genetic information processing (i.e. replication and repair; translation; folding sorting and degradation) and nucleotide metabolism in gut bacterial communities can namely be explained by the importance of DNA repair systems for bacterial colonization of intestines (Vogel-Scheel et al. 2010, Davies et al. 2011). Reactive nitrogen species and oxidative stress indeed play an important role in host defense against pathogens and have been documented to regulate microbiome assembly in animal hosts (O’Rourke et al. 2003, Fang and Vasquez-Torres 2020).

A functional cluster composed of pathways related to xenobiotics degradation and metabolism and to amino acid metabolism were comparatively important in explaining bacterial community composition in soil and plant microbiota. The prevalence of these metabolic functions can be linked to the variety of aromatic compounds (i.e. xenobiotics) and amino acids that plant tissues contain and exude in the soil, which represent a common and varied source of carbon for terrestrial bacteria (Moe 2013, Pérez-Pantoja et al. 2015). The ability to degrade plant aromatic compounds was for example linked to a better colonization and growth of bacteria in rhizosphere habitats (Ledger et al. 2012). Degradation of antibiotic compounds secreted by other microbes and macro-organisms are also important for survival in the dense and diverse biotic networks that interact in the soil (Lucas et al. 2019).

Functional pathways linked with the conversion of light to sugar (i.e. photosynthesis and the TCA cycle), and with the metabolism of cofactors and vitamins were predominant in bacterial communities found in aquatic habitats, and especially in saltwater, outlining the importance of autotrophy for bacterial life in aquatic ecosystems. Light-driven proton pumps (i.e. proteorhodopsins) are for example used by a variety of *Flavobacteriia* to generate energy in high-light marine environments (Kumagai et al. 2018). The enrichment of these types of functions in aquatic habitats was contrasted with the enrichment of cell motility pathways in terrestrial habitats, which could be explained by a more general role for bacterial motility in tracking nutrients and carbon sources in the soil (i.e. for heterotrophic lifestyles - Turnbull et al. 2001), or in engaging with plants and animals (Akahoshi and Bevins 2022). Finally, the prevalence of cofactors and vitamin metabolic pathways in saltwater bacteria can be linked to the documented depletion of several vitamins in oceanic habitats, which has been proposed as an important filter for planktonic community assembly (Sañudo-Wilhelmy et al. 2012).

Our analyses also identified clusters that explained variation between endophytic and bacteria and other host-associated bacteria. These clusters both contained functions linked with community sensing and cell-cell interaction, which likely facilitate establishment on, and interaction with the host. Functional pathways involved in biofilms were more present in non-endophytic bacteria, which can be linked to their role in facilitating colonization of host external and internal surfaces (Bogino et al. 2013, Motta et al. 2021). Quorum sensing functions were comparatively enriched in endophytic communities where they can contribute to bacterial entry in the host tissues (Gosai et al. 2020).

The functional classification of bacterial communities portrayed here among habitats is similar in some ways to those suggested in other studies of microbial functional variation. Dinsdale et al. (2008) identified functions associated with certain habitats, but focused on broad, high-level functional categories. Other studies (e.g. Ramírez-Flandes et al. 2019, Malik et al. 2020) found that different genes allowed the functional classification of microbial communities into different habitat types but focused on a more limited range of habitat types (for example, on soil microbial communities). By expanding analyses of bacterial functions to identify functional strategies across metagenomes and genomes without an a priori selection of subsets of functions or habitats to examine, our analyses revealed that major axes of bacterial trait variation are structured by clusters of correlated traits that correspond to coherent functional categories across diverse habitats. It remains an open question whether ecological strategies identified for macroorganisms such as plants are relevant for understanding the strategies of microorganisms. In this study, we did not find clear mappings between major groups of correlated traits explaining variation among bacterial genomes and habitats and the competitor-stress tolerator-ruderal (CSR) scheme of Grime (1977), whose application to microbial communities has recently been proposed (Fierer 2017). Our results overall emphasize the role of data-driven approaches in providing novel insights into the ecological and evolutionary factors that drive microbial trait variation.

### Phylogenetic structure of functional pathways and clusters

Phylogenetically constrained functional clusters based on genomic data were generally correlated with those obtained from metagenomic data. Concordantly, clusters of functions that were associated with the first axis of the PCoA and thus explained the most functional variation among habitats (C2, C9) were also the most phylogenetically ‘conserved’, having the strongest signal in the genomic phylogeny across most depths. These results suggest that selection or constraint (e.g. genetic correlations (Futuyma 2010) or functional barriers to recombination (Hendrickson et al. 2018)) have been important in shaping the evolution of clusters of correlated traits in bacteria in a consistent fashion across both clades and habitats, leading to predictable variation in the ecological strategies of bacteria associated with particular clades. The fact that we found such clusters despite the likely presence of many transient organisms in environmental metagenomes provides additional confidence in these results.

Despite identifying three main axes of functional covariation among bacterial traits, other drivers of functional trait correlations appeared to play a role in structuring trait variation among ecosystems (Appendix S1: Table S1). First, some metagenomically-derived trait clusters (e.g. C4, C5, C8) were not observed in the genomic dataset, suggesting that they might result from environmental filtering on pre-existing trait variation or lateral gene transfer among host-associated clades rather than a fundamental constraint or trade-off in genome architecture and evolution. These clusters still showed some level of phylogenetic signal in the genomic phylogeny, but more so at shallower phylogenetic depths, consistent with filtering of genes transferred horizontally among related organisms living in a shared habitat (Shapiro et al. 2012). Second, traits belonging to the cluster C7 did not explain much variation across the axes and were not enriched in any particular type of high-level functional categories, suggesting they do not represent part of a functional strategy that has arisen in response to the major axes of environmental variation encompassed by this dataset. Their lack of correlations in the genomic dataset as well suggests that they are likely to have been evolving idiosyncratically or to be subject to more extensive horizontal gene transfer. It is possible that other types of functional strategies than those identified above are also important for driving within-community niche partitioning among bacteria, but at the broad scales of the bacterial tree of life and among metagenomes from distinct habitat types, these types of trade-offs are not important enough to be consistently identified as clusters of covarying traits. Testing how important the main functional axes identified in this study are in explaining microbial fitness at smaller spatial and environmental scales will represent a valuable next research direction for the field.

## Conclusion

In conclusion, we used a data-driven functional trait screening approach to identify the major axes of functional trait covariation in bacterial metagenomes and genomes based on gene functional annotations. By comparing functional trait clusters in both datasets, we show that the major strategies driving functional differentiation between metagenomes of co-occurring bacteria in ecological communities tend to be phylogenetically structured among bacterial genomes. We also point to the limited overlap between the bacterial ecological strategy axes described here and the trait classification schemes developed for plants and animals. By reducing the high-dimensionality of trait variation observed among microorganisms around a small number of fundamental axes of trait covariation, we make an important step towards generalization of microbial ecology and of the drivers of biological diversity and bacterial biogeography across study systems.

## Data availability statement

All metagenomic datasets used in this study can be accessed either on the IMG/M (https://img.jgi.doe.gov/cgi-bin/m/main.cgi) or the MG-RAST (https://www.mg-rast.org/) online databases using the sample identification information provided in Appendix S1: Table S2. The functional and phylogenetic genomic datasets can be accessed within the AnnoTree database (http://annotree.uwaterloo.ca/app/downloads.html) using the sample identification information provided in Appendix S1: Table S3. The novel R code developed for processing and analyzing the data is available for review through Figshare (https://figshare.com/s/2d9835e083f160939d66) and will be made public in a GitHub repository (https://github.com/glajoie1/Bact_Fun_Axes) upon publication of the manuscript.

## Acknowledgments

We are grateful to Jesse Shapiro and Mario Muscarella for comments on a version of this manuscript. We acknowledge funding from the Natural Sciences and Engineering Research Council of Canada (GL, SWK) and the Canada Research Chairs (SWK).

## Literature cited

Ackerly, D. D., and P. B. Reich. 1999. Convergence and correlations among leaf size and function in seed plants: comparative tests using independent contrast. American Journal of Botany 86:1272–1281.

Akahoshi, D. T., and C. L. Bevins. 2022. Flagella at the host-microbe interface: Key functions intersect with redundant responses. Frontiers in Immunology 13:828758.

Bogino, P. C., M. de la Mercedes Oliva, F. Sorroche, and W. Giordano. 2013. The role of bacterial biofilms and surface components in plant-bacterial associations. International Journal of Molecular Sciences 14:15838–15859.

Charrad, M., N. Ghazzali, V. Boiteau, and A. Niknafs. 2014. NbClust: An R Package for determining the relevant number of clusters in a data set. Journal of Statistical Software 61:1–36.

Chen, I. A., K. Chu, K. Palaniappan, M. Pillay, A. Ratner, J. Huang, M. Huntemann, N. Varghese, J. R. White, R. Seshadri, T. Smirnova, E. Kirton, S. P. Jungbluth, T. Woyke, N. N. Ivanova, and N. C. Kyrpides. 2019. IMG/M v.5.0: an integrated data management and comparative analysis system for microbial genomes and microbiomes. Nucleic Acids Research 47:666–677.

Clooney, A. G., F. Fouhy, R. D. Sleator, and A. O. Driscoll. 2016. Comparing apples and oranges ?: Next generation sequencing and its impact on microbiome analysis. PLoS ONE 11:e0148028.

Davies, B. W., R. W. Bogard, N. M. Dupes, T. A. I. Gerstenfeld, L. A. Simmons, and J. John. 2011. DNA damage and reactive nitrogen species are barriers to *Vibrio cholerae* colonization of the infant mouse intestine. PLoS Pathogens 7:e1001295.

Díaz, S., J. Kattge, J. H. C. Cornelissen, I. J. Wright, S. Lavorel, S. Dray, B. Reu, M. Kleyer, C. Wirth, I. C. Prentice, E. Garnier, G. Bönisch, M. Westoby, H. Poorter, P. B. Reich, A. T. Moles, J. Dickie, A. N. Gillison, A. E. Zanne, J. Chave, S. J. Wright, S. N. Sheremet’ev, H. Jactel, B. Christopher, B. Cerabolini, S. Pierce, B. Shipley, D. Kirkup, F. Casanoves, J. S. Joswig, A. Günther, V. Falczuk, N. Rüger, M. D. Mahecha, and L. D. Gorné. 2016. The global spectrum of plant form and function. Nature 529:167–171.

Dinsdale, E. A., R. A. Edwards, D. Hall, F. Angly, M. Breitbart, J. M. Brulc, M. Furlan, C. Desnues, M. Haynes, L. Li, L. McDaniel, M. A. Moran, K. E. Nelson, C. Nilsson, R. Olson, J. Paul, B. R. Brito, Y. Ruan, B. K. Swan, R. Stevens, D. L. Valentine, R. V. Thurber, L. Wegley, B. A. White, and F. Rohwer. 2008. Functional metagenomic profiling of nine biomes. Nature 452:629–632.

Evans, S. E., and M. D. Wallenstein. 2014. Climate change alters ecological strategies of soil bacteria. Ecology Letters 17:155–164.

Fang, F. C., and A. Vasquez-Torres. 2020. Reactive nitrogen species in host-bacterial interactions. Current Opinion in Immunology 60:96–102.

Fierer, N. 2017. Embracing the unknown: disentangling the complexities of the soil microbiome. Nature Reviews Microbiology 15:579–590.

Fierer, N., M. A. Bradford, and R. B. Jackson. 2007. Toward an ecological classification of soil bacteria. Ecology 88:1354–1364.

Fierer, N., J. W. Leff, B. J. Adams, U. N. Nielsen, S. Thomas, C. L. Lauber, S. Owens, J. A. Gilbert, D. H. Wall, and J. G. Caporaso. 2012. Cross-biome metagenomic analyses of soil microbial communities and their functional attributes. PNAS 109:21390–21395.

Futuyma, D. J. 2010. Evolutionary constraint and ecological consequences. Evolution 64:1865–1884.

Galili, T. 2015. dendextend: an R package for visualizing, adjusting, and comparing trees of hierarchical clustering. Bioinformatics 31:3718–3720.

Genz, A., F. Bretz, T. Miwa, X. Mi, F. Leisch, F. Scheipl, and T. Hothorn. 2021. mvtnorm: Multivariate normal and t distributions. R Package version 1.1-3.

Gosai, J., S. Anandhan, A. Bhattacharjee, and G. Archana. 2020. Elucidation of quorum sensing components and their role in regulation of symbiotically important traits in Ensifer nodulating pigeon pea. Microbiological Research 231:126354.

Grime, J. P. 1977. Evidence for the existence of three primary strategies in plants and its relevance to ecological and evolutionary theory. The American Naturalist 111:1169–1194.

Hendrickson, H. L., D. Barbeau, R. Ceschin, and J. G. Lawrence. 2018. Chromosome architecture constrains horizontal gene transfer in bacteria. PLoS Genetics 14:e1007421.

Jombart, T., and S. Dray. 2010. adephylo: exploratory analyses for the phylogenetic comparative method. Bioinformatics 26:1907–1909.

Jombart, T., S. Pavoine, S. Devillard, and D. Pontier. 2010. Putting phylogeny into the analysis of biological traits: a methodological approach. Journal of Theoretical Biology 264:693–701.

Kanehisa, M., S. Goto, Y. Sato, M. Kawashima, M. Furumichi, and M. Tanabe. 2014. Data, information, knowledge and principle: Back to metabolism in KEGG. Nucleic Acids Research 42:199–205.

Keck, F., F. Rimet, A. Bouchez, and A. Franc. 2016. phylosignal: an R package to measure, test, and explore the phylogenetic signal. Ecology and Evolution 6:2774–2780.

Keegan, K., E. Glass, and F. Meyer. 2016. MG-RAST, a metagenomics service for analysis of microbial community structure and function. Methods in Molecular Biology 1399:207–233.

Kumagai, Y., S. Yoshizawa, Y. Nakajima, M. Watanabe, T. Fukunaga, Y. Ogura, T. Hayashi, K. Oshima, M. Hattori, and M. Ikeuchi. 2018. Solar-panel and parasol strategies shape the proteorhodopsin distribution pattern in marine *Flavobacteriia*. The ISME Journal 12:1329–1343.

Lajoie, G., and S. W. Kembel. 2019. Making the most of trait-based approaches for microbial ecology. Trends in Microbiology 27:814–823.

Leary, N. A. O., M. W. Wright, J. R. Brister, S. Ciufo, D. Haddad, R. Mcveigh, B. Rajput, B. Robbertse, B. Smith-white, D. Ako-adjei, A. Astashyn, A. Badretdin, Y. Bao, O. Blinkova, V. Brover, V. Chetvernin, J. Choi, E. Cox, O. Ermolaeva, C. M. Farrell, T. Goldfarb, T. Gupta, D. Haft, E. Hatcher, W. Hlavina, S. Joardar, V. K. Kodali, W. Li, D. Maglott, P. Masterson, M. Mcgarvey, M. R. Murphy, K. O. Neill, S. Pujar, S. H. Rangwala, D. Rausch, L. D. Riddick, C. Schoch, A. Shkeda, S. S. Storz, H. Sun, F. Thibaud-nissen, I. Tolstoy, R. E. Tully, R. Vatsan, C. Wallin, D. Webb, W. Wu, M. J. Landrum, A. Kimchi, T. Tatusova, M. Dicuccio, P. Kitts, T. D. Murphy, and K. D. Pruitt. 2016. Reference sequence (RefSeq) database at NCBI: current status, taxonomic expansion, and functional annotation. Nucleic Acids Research 44:733–745.

Ledger, T., A. Zuniga, T. Kraiser, P. Dasencich, R. Donoso, D. Pérez-Pantoja, and B. Gonzalez. 2012. Aromatic compounds degradation plays a role in colonization of *Arabidopsis thaliana* and *Acacia caven* by *Cupriavidus pinatubonensis* JMP134. Antonie van Leeuwenhoek 101:713–723.

Lenth, R. 2022. Package ‘Estimated Marginal Means, aka Least-Squares Means’.

Lucas, J. M., E. Gora, A. Salzberg, M. Kaspari, and J. M. Lucas. 2019. Antibiotics as chemical warfare across multiple taxonomic domains and trophic levels in brown food webs. Proceedings of the Royal Society B: Biological Sciences.

Madani, N., J. S. Kimball, A. P. Ballantyne, D. L. R. Affleck, P. M. Van Bodegom, P. B. Reich, J. Kattge, A. Sala, M. Nazeri, M. O. Jones, M. Zhao, and S. W. Running. 2018. Future global productivity will be affected by plant trait response to climate. Scientific Reports 8:2870.

Malik, A. A., J. B. H. Martiny, E. L. Brodie, A. C. Martiny, K. K. Treseder, and S. D. Allison. 2020. Defining trait-based microbial strategies with consequences for soil carbon cycling under climate change. The ISME Journal 14:1–9.

Mendler, K., H. Chen, D. H. Parks, B. Lobb, L. A. Hug, and A. C. Doxey. 2019. AnnoTree: visualization and exploration of a functionally annotated microbial tree of life. Nucleic Acids Research 47:4442–4448.

Moe, L. 2013. Amino acids in the rhizosphere: from plants to microbes. American Journal of Botany 100:1692–1705.

Motta, J., J. Wallace, A. Buret, C. Deraison, and N. Vergnolle. 2021. Gastrointestinal biofilms in health and disease. Nature Reviews Gastroenterology & Hepatology 18:314–334.

Muir, C. D. 2015. Making pore choices: repeated regime shifts in stomatal ratio. Proceedings of the Royal Society B: Biological Sciences 282:20151498.

O’Rourke, E. J., C. Chevalier, A. V. Pinto, J. M. Thiberge, L. Ielpi, and J. P. Radicella. 2003. Pathogen DNA as target for host-generated oxidative stress: Role for repair of bacterial DNA damage in *Helicobacter pylori* colonization. PNAS. 100:2789–2794.

Oksanen, A. J., F. G. Blanchet, M. Friendly, R. Kindt, P. Legendre, D. Mcglinn, P. R. Minchin, R. B. O. Hara, G. L. Simpson, P. Solymos, M. H. H. Stevens, and E. Szoecs. 2013. Package ‘ vegan’

Paradis, E., and K. Schliep. 2018. ape 5.0: an environment for modern phylogenetics and evolutionary analyses in R. Bioinformatics 35:526–528.

Pérez-Pantoja, D., P. Leiva-Novoa, R. A. Donoso, C. Little, M. Godoy, D. H. Pieper, and B. Gonzàlez. 2015. Hierarchy of carbon source utilization in soil bacteria: Hegemonic preference for benzoate in complex aromatic compound mixtures degraded by *Cupriavidus pinatubonensis* strain JMP134. Applied and Environmental Microbiology 81:3914–3924.

Ramírez-Flandes, S., B. González, and O. Ulloa. 2019. Redox traits characterize the organization of global microbial communities. PNAS 116:3630–3635.

Santillan, E., H. Seshan, F. Constancias, and S. Wuertz. 2019. Trait-based life-history strategies explain succession scenario for complex bacterial communities under varying disturbance. Environmental Microbiology 21:3751–3764.

Sañudo-Wilhelmy, S. A., L. S. Cutter, R. Durazo, E. A. Smail, L. Gómez-Consarnau, E. Webb, M. Prokopenko, W. Berelson, and D. Karl. 2012. Multiple B-vitamin depletion in large areas of the coastal ocean. PNAS 109:14041–14045.

Shapiro, B., J. Friedman, O. Cordero, S. Preheim, S. Timberlake, G. Szabó, M. F. Polz, and E. J. Alm. 2012. Population genomics of early events in the ecological differentiation of bacteria. Science 336:48–51.

Steen, A. D., A. C. P. Carini, K. G. Lloyd, J. C. Thrash, K. M. Deangelis, and N. Fierer. 2019. High proportions of bacteria and archaea across most biomes remain uncultured. The ISME Journal:3126–3130.

Turnbull, G. A., J. A. W. Morgan, J. M. Whipps, and J. R. Saunders. 2001. The role of bacterial motility in the survival and spread of *Pseudomonas fluorescens* in soil and in the attachment and colonisation of wheat roots. FEMS Microbiology Ecology 36:21–31.

Vogel-Scheel, J., C. Alpert, W. Engst, G. Loh, M. Blaut, and H. N. Potsdam-rehbru. 2010. Requirement of purine and pyrimidine synthesis for colonization of the mouse intestine by Escherichia coli. Applied and Environmental Microbiology 76:5181–5187.

Weisberg, F. 2019. An R Companion to Applied Regression. Sage, Thousand Oaks, California.

Westoby, M. 1998. A leaf-height-seed (LHS) plant ecology strategy scheme. Plant and Soil 199:213–227.

Wright, I. J., P. B. Reich, M. Westoby, D. D. Ackerly, Z. Baruch, F. Bongers, J. Cavender-bares, T. Chapin, J. H. C. Cornelissen, M. Diemer, J. Flexas, E. Garnier, P. K. Groom, and J. Gulias. 2004. The worldwide leaf economics spectrum. Nature 428:821–827.

