## Appendix S1 for "Data-driven identification of major axes of functional variation in bacteria"

### Data-driven identification of major axes of functional variation in bacteria

Geneviève Lajoie & Steven W. Kembel

*Environmental Microbiology*

**Table S1. Predictions of the ecological and evolutionary significance of groups of functional pathways based on their clustering in the metagenomic and the genomic datasets.**

**Table S2. Metadata of metagenomic samples used in this study.** The ID.Label column represents sample identifier used in Figure 1.

**Table S3. Accession numbers of the genomes used in this study from the Genome Taxonomy Database (GTDB).**

**Table S4. Functional pathways composing every metagenomic cluster and their functional categorization in the KEGG BRITE hierarchy.**

**Table S5. Type-III analysis of variance (ANOVA) of the phylogenetic signal (Moran's I) of pathways in the genomic phylogeny as a function of the cluster in which they are found, the phylogenetic depth at which the signal was calculated and their interaction.**

**Table S6. Analyses of variance of the phylogenetic signal (Moran's I) of functional pathways in the genomic phylogeny as a function of the correlation of these pathways with each of the first four dimension of the metagenomic PCoA (Figure 2), the phylogenetic depth at which the signal was calculated, and the interaction between these factors.** Models were conducted separately per PCoA dimension and direction along this dimension (positive or negative).

**Table S1.** Predictions of the ecological and evolutionary significance of groups of functional pathways based on their clustering in the metagenomic and the genomic datasets.

| | | **Genomic dataset** | |
|---|---|---|---|
| | **Group of traits is :** | **Clustered** | **Not clustered** |
| **Metagenomic dataset** | **Clustered** | The cluster is a conserved strategy of current ecological importance. (There is selection and/or constraints on the evolution of this group of traits.) | The cluster represents a strategy selected in the species pool, regardless of phylogenetic identity. |
| | **Not clustered** | The cluster may have been selected for under previous ecological constraints that are no longer important in driving the distribution of organisms among ecosystems, or that explain ecological variation at another scale. | These traits do not participate in a general strategy.<br>They might be labile traits that do not have strong constraints of evolution.<br>They might be important for occupying rare niches. |

Appendix S1

Data-driven identification of major axes of bacterial functional variation

Geneviève Lajoie & Steven W. Kembel

*Environmental Microbiology*

**Table S1. Predictions of the ecological and evolutionary significance of groups of functional pathways based on their clustering in the metagenomic and the genomic datasets.**

**Table S2. Metadata of metagenomic samples used in this study.** The ID.Label column represents sample identifier used in Figure 1.

**Table S3. Accession numbers of the genomes used in this study from the Genome Taxonomy Database (GTDB).**

**Table S4. Functional pathways composing every metagenomic cluster and their functional categorization in the KEGG BRITE hierarchy.**

**Table S5. Type-III analysis of variance (ANOVA) of the phylogenetic signal (Moran's I) of pathways in the genomic phylogeny as a function of the cluster in which they are found, the phylogenetic depth at which the signal was calculated and their interaction.**

**Table S6. Analyses of variance of the phylogenetic signal (Moran's I) of functional pathways in the genomic phylogeny as a function of the correlation of these pathways with each of the first four dimension of the metagenomic PCoA (Figure 2), the phylogenetic depth at which the signal was calculated, and the interaction between these factors.** Models were conducted separately per PCoA dimension and direction along this dimension (positive or negative).

**Table S1.** Predictions of the ecological and evolutionary significance of groups of functional pathways based on their clustering in the metagenomic and the genomic datasets.

| | | **Genomic dataset** | |
|---|---|---|---|
| | **Group of traits is :** | **Clustered** | **Not clustered** |
| **Metagenomic dataset** | **Clustered** | The cluster is a conserved strategy of current ecological importance. (There is selection and/or constraints on the evolution of this group of traits.) | The cluster represents a strategy selected in the species pool, regardless of phylogenetic identity. |
| | **Not clustered** | The cluster may have been selected for under previous ecological constraints that are no longer important in driving the distribution of organisms among ecosystems, or that explain ecological variation at another scale. | These traits do not participate in a general strategy.<br>They might be labile traits that do not have strong constraints of evolution.<br>They might be important for occupying rare niches. |
