## Appendix S2 for "Data-driven identification of major axes of functional variation in bacteria"

### Data-driven identification of major axes of functional variation in bacteria

Geneviève Lajoie & Steven W. Kembel

*Environmental Microbiology*

**Figure S1. Rarefaction curves for the number of functional genes (panel a) and the number of taxonomic families (panel b) per metagenomic sample.** These curves were generated after filtering out sequences that could not be both functionally and taxonomically annotated. Each curve represents a metagenomic sample. For the figure, sample sizes were cut-off at a limit of 2 000 000 to facilitate visualization of the curvature points of the curves. Samples that had more annotated sequences than this threshold plateaued beyond that point such that the interpretation of these curves is not affected by that procedure.

**Figure S2. Taxonomic composition of bacterial genomes and metagenomic samples included in this study.** Mean relative frequencies of sequences from each phylum were calculated across samples for each habitat. Phyla that had relative abundances lower than 0.5% were grouped together in a low-frequency phyla category.

**Figure S3. Detailed composition of each cluster obtained in the hierarchical clustering of bacterial functional pathways across metagenomic samples.** The Tier 2 KEGG functional category to which each pathway belongs are identified by the color and shape of the symbol (see legend). The order of functional pathways (from top to bottom) in each cluster corresponds to that presented (from left to right) in Figure 2.

**Figure S4. Richness and Shannon diversity of functional pathways in metagenomic samples from different habitats.** Letters above the boxplots indicate whether the mean diversity of two given habitats is significantly different (their letters are different) or not (they share at least one letter). Differences among habitats were tested for each diversity index using a type-III ANOVA with post-hoc pairwise comparisons between habitats calculated using estimated marginal means. Pairwise comparisons were adjusted for multiple testing using the multivariate t distribution method (Genz et al. 2021).

**Figure S5. Variation in the composition of functional pathways among metagenomic samples that can be explained as a function of habitat (left), taxonomic class (right) and either factor (intersect).**

**Figure S6. Phylogenetic signal (Moran's *I*) of functional pathways across functional categories and phylogenetic depths.** Moran's *I* of each functional pathway were calculated using phylogenetic correlograms across the bacterial genomic phylogeny, from the tip (depth = 0) towards the root of the tree (depth = 0.0444). Black dots represent the value of estimated marginal means of Moran's I across the functional pathways of each

Tier I KEGG functional category, at each of five depths. The blue bars are confidence intervals. The absence of overlap among red arrows of two clusters at a given depth indicates that their mean phylogenetic signals are significantly different. These pairwise comparisons were adjusted for multiple testing using the multivariate *t* distribution method (Genz et al. 2021).

**Figure S7. Relationship between the phylogenetic signal (Moran's I) of functional pathways in the genomic phylogeny and their contribution to variation along the first four dimensions of the principal coordinates analysis performed on the functional composition of metagenomic samples** (see Figure 1). Points represent individual functional pathways. Regressions were calculated separately for pathways that had positive and negative correlations with a given dimension, and in each case across 5 phylogenetic depths. Moran's *I* of each functional pathway were calculated using phylogenetic correlograms across the bacterial genomic phylogeny, from the tip (depth = 0) towards the root of the tree (depth = 0.0444). Slopes are shown when statistically significant.

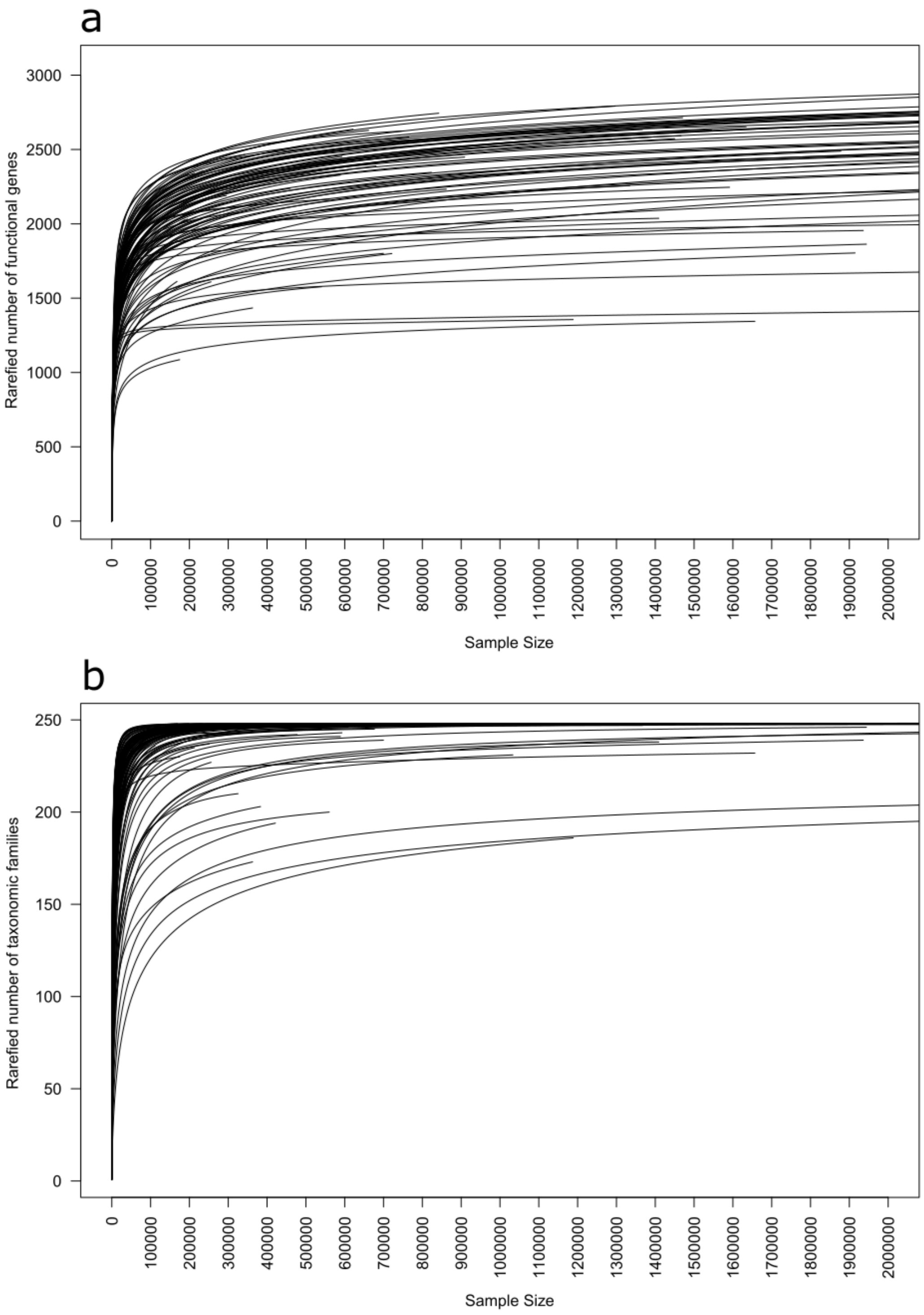
a
Rarefied number of functional genes
0
500
1000
1500
2000
2500
3000
Sample Size
0
100000
200000
300000
400000
500000
600000
700000
800000
900000
1000000
1100000
1200000
1300000
1400000
1500000
1600000
1700000
1800000
1900000
2000000
b
Rarefied number of taxonomic families
0
50
100
150
200
250
Sample Size

Figure S1.

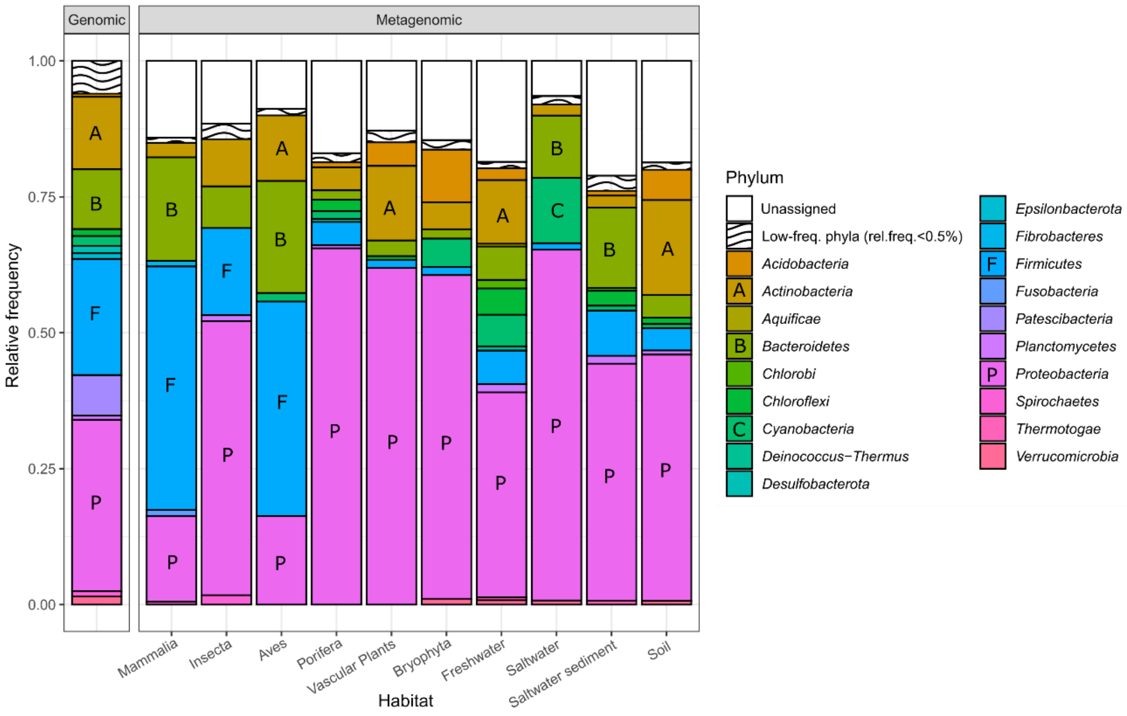
Genomic
Metagenomic
Relative frequency
1.00
0.75
0.50
0.25
0.00
A
B
F
P
Mammalia
Insecta
Aves
Porifera
Vascular Plants
Bryophyta
Freshwater
Saltwater
Saltwater sediment
Soil
Habitat
C
Phylum
Unassigned
Low-freq. phyla (rel.freq.<0.5%)
Acidobacteria
Actinobacteria
Aquificae
Bacteroidetes
Chlorobi
Chloroflexi
Cyanobacteria
Deinococcus−Thermus
Desulfobacterota
Epsilonbacterota
Fibrobacteres
Firmicutes
Fusobacteria
Patescibacteria
Planctomycetes
Proteobacteria
Spirochaetes
Thermotogae
Verrucomicrobia

Figure S2.

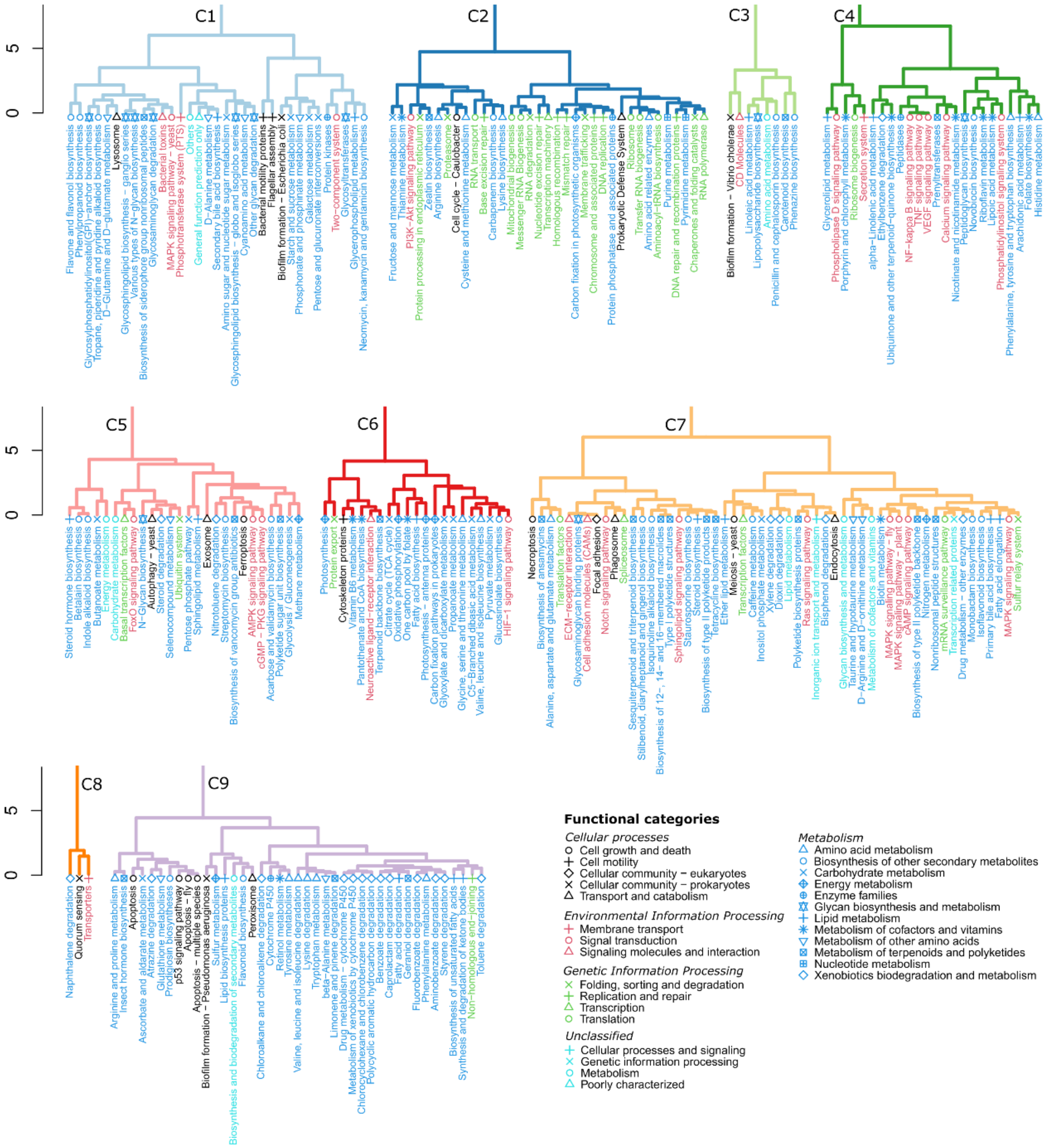
C1
C2
C3
C4
C5
C6
C7
C8
C9
0
5
Functional categories
Cellular processes
Cell growth and death
Cell motility
Cellular community − eukaryotes
Cellular community − prokaryotes
Transport and catabolism
Environmental Information Processing
Membrane transport
Signal transduction
Signaling molecules and interaction
Genetic Information Processing
Folding, sorting and degradation
Replication and repair
Transcription
Translation
Unclassified
Cellular processes and signaling
Genetic information processing
Metabolism
Poorly characterized
Metabolism
Amino acid metabolism
Biosynthesis of other secondary metabolites
Carbohydrate metabolism
Energy metabolism
Enzyme families
Glycan biosynthesis and metabolism
Lipid metabolism
Metabolism of cofactors and vitamins
Metabolism of other amino acids
Metabolism of terpenoids and polyketides
Nucleotide metabolism
Xenobiotics biodegradation and metabolism

Figure S3.

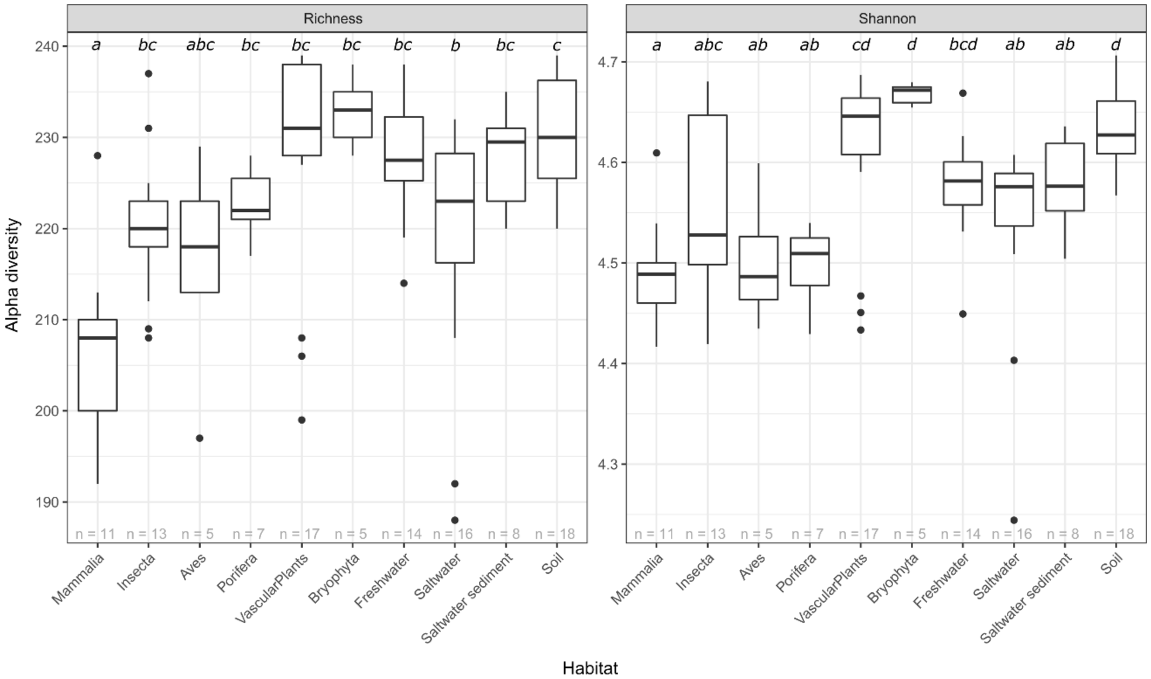

Richness
Shannon
Alpha diversity
a bc abc bc bc bc bc b bc c
a abc ab ab cd d bcd ab ab d
240
230
220
210
200
190
4.7
4.6
4.5
4.4
4.3
n = 11 n = 13 n = 5 n = 7 n = 17 n = 5 n = 14 n = 16 n = 8 n = 18
Mammalia
Insecta
Aves
Porifera
VascularPlants
Bryophyta
Freshwater
Saltwater
Saltwater sediment
Soil
Habitat


Figure S4.

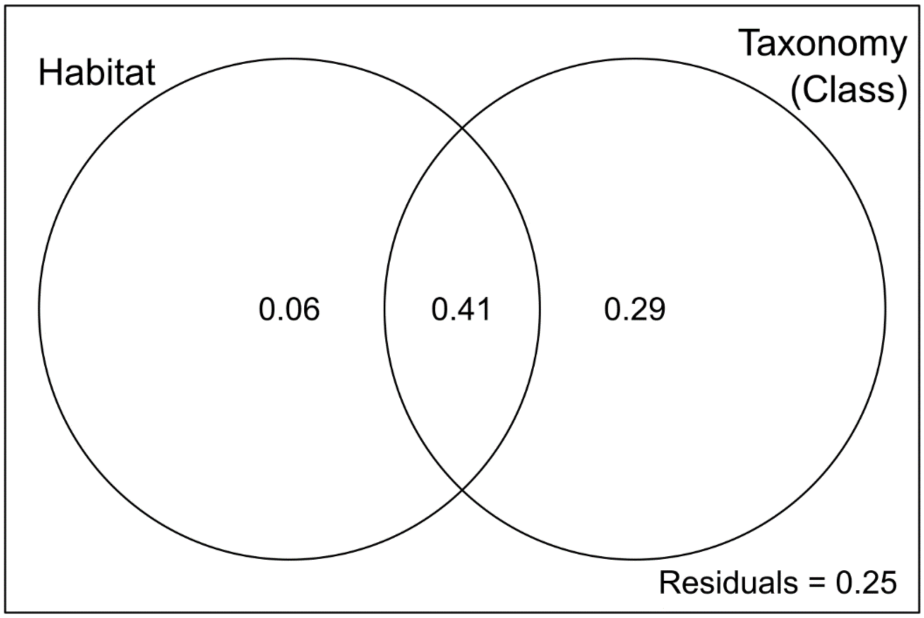


Figure S5.

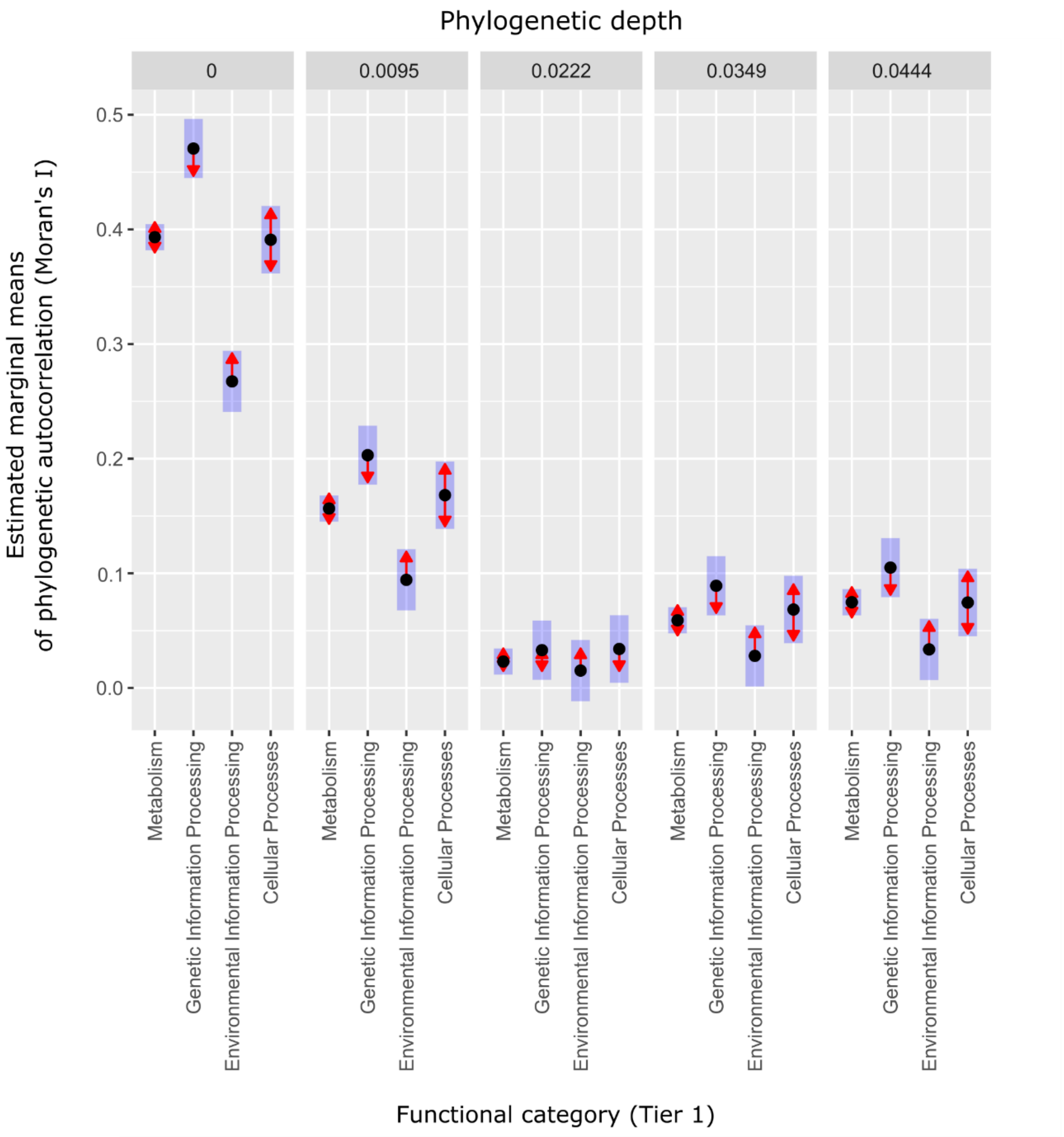

Phylogenetic depth
0
0.0095
0.0222
0.0349
0.0444
Estimated marginal means
of phylogenetic autocorrelation (Moran's I)
0.5
0.4
0.3
0.2
0.1
0.0
Metabolism
Genetic Information Processing
Environmental Information Processing
Cellular Processes
Functional category (Tier 1)


Figure S6.

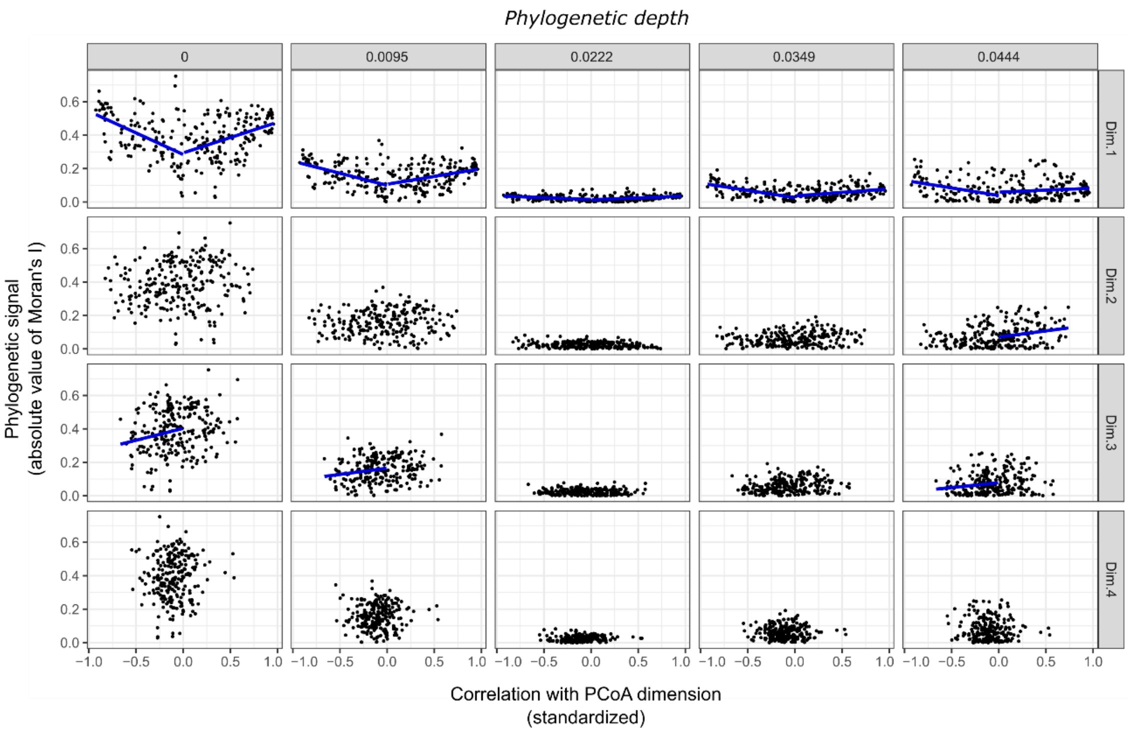

Phylogenetic depth
0
0.0095
0.0222
0.0349
0.0444
Dim.1
Dim.2
Dim.3
Dim.4
Phylogenetic signal
(absolute value of Moran's I)
Correlation with PCoA dimension
(standardized)


Figure S7.

#### References

Genz, A., F. Bretz, T. Miwa, X. Mi, F. Leisch, F. Scheipl, and T. Hothorn. 2021. mvtnorm: Multivariate normal and t distributions. R Package version 1.1-3.
